# Network analysis of canine brain morphometry links tumour risk to oestrogen deficiency and accelerated brain ageing

**DOI:** 10.1101/412643

**Authors:** Nina M. Rzechorzek, Olivia M. Saunders, Lucy Hisco, Tobias Schwarz, Katia Marioni-Henry, David J. Argyle, Jeffery J. Schoenebeck, Tom C. Freeman

## Abstract

**Background:** Structural ‘brain age’ is a valuable but complex biomarker for several brain disorders. The dog is an unrivalled comparator for neurological disease modeling, however brain phenotypic diversity among pedigrees creates computational and statistical challenges.

**Methods:** We applied unbiased network correlation analysis in dogs to explore complex interactions between brain morphometrics, patient metadata, and neurological disease. Twenty-four parameters measured from each of 286 brain magnetic resonance imaging scans generated 9,438 data points that were used to cluster canine patients according to their brain morphometry profiles. The network was then explored for statistically significant enrichments within breed, sex, age, and diagnostic categories.

**Findings:** Morphometric comparisons revealed an advanced ‘aged-brain’ profile in the Boxer breed, consisting of a small brain length, width, and volume, combined with ventriculomegaly. Key features of this profile were paralleled in neutered female dogs which, relative to un-neutered females, had an 11-fold greater risk of developing primary brain tumours. Enrichment analysis confirmed that Boxers and geriatric individuals were enriched for brain tumour diagnoses, despite a lack of geriatric Boxers within the cohort.

**Interpretation:** These findings suggest that accelerated brain ageing might contribute to tumour risk in Boxers and may be influenced by oestrogen deficiency — a risk factor for dementia and brain tumours in humans. We propose that morphometric features of brain ageing in dogs, like humans, might better predict neurological disease risk than a patient’s chronological age.

**Funding:** Wellcome Trust Integrated Training Fellowship for Veterinarians (096409/Z/11/Z to N.M.R) and an MSD Animal Health Connect Bursary (to O.M.S.).

## Research in context

### Evidence before this study

Our ageing human population has increased the prevalence of chronic brain disorders, placing an unsustainable burden on healthcare systems worldwide. Contributing to this problem is our limited understanding of how chronic brain disorders begin, how they might be treated, and at what disease stage treatment is most likely to benefit patients. The domestic dog is fast becoming the most important species for neurological disease modeling; dogs naturally develop many of the same brain disorders as humans, and their highly controlled genetics, shorter lifespan, and shared environment with us makes them ideal ‘companions’ for comparative brain studies. It is well recognized that brain ageing varies among individuals, and that our biological or structural ‘brain age’ better predicts our risk of brain disease than our age in years. This discrepancy may be exaggerated in the domestic dog, where selective breeding has produced an extreme diversity in age-related brain changes and brain shape across pedigrees. If we can better understand how brain shape varies with age and disease, we can use aspects of brain shape to predict which patients would most benefit from early intervention.

Certain brain features are expected to change as we age, but these changes are subtle, complex, and may affect some parts of the brain more than others. Sophisticated machine learning techniques can be used to estimate biological ‘brain age’ from human brain scans, but such techniques depend on the fact that age-related changes in brain structure are very similar among healthy humans. By contrast, dramatic variations in head shape between different dog breeds makes automated analyses very difficult, and there is limited published data describing the structural diversity of the canine brain. In many countries, magnetic resonance imaging (MRI) facilities are now accessible to canine patients, and MRI has become an indispensible tool for veterinary neurologists. Canine patient datasets provide a rich, but complex picture of disease on a background of both individual- and breed-based variation. Here we have used an unbiased, data-driven method called network analysis to make sense of this complexity. Our objective was to understand how brain shape and other canine patient factors impact on neurological disease risk.

### Added value of this study

With an unconventional data-driven approach, we have identified an advanced ‘brain age’ in neutered female dogs and the Boxer breed that is associated with brain tumour risk. To our knowledge, this is the first study to link brain tumour risk to both accelerated brain ageing and oestrogen loss. Moreover, we have demonstrated that neutering status affects brain shape in dogs. This study outlines new collaborative opportunities — through comparative biology — to understand the influence of chromosomal and hormonal sex on brain structure, brain ageing, and brain tumour development. Our data can also be incorporated into larger canine databases to help extract non-invasive image-based biomarkers for neurological disease. Furthermore, our unique network analysis approach to handle the combined complexity within clinical and brain imaging datasets is immediately relevant to human patient studies.

### Implications of all the available evidence

Extreme breed characteristics impact on animal health and welfare, with current widespread concerns over dogs with ‘short faces’. Our work extends this to the brain, highlighting an urgency to better understand factors that influence brain ageing in dogs. Further studies are needed to confirm how patient, lifestyle, and environmental factors influence brain structure throughout the life course. Our approach can help reveal subtle yet important changes within these complex datasets. We propose that the role of sex in brain ageing would be more readily understood by studying dogs of varied neutering status, not least because neutering is a non-obligatory but common practice that routinely takes place at a defined, early life-stage. In addition, our results identify the Boxer breed as a potentially valuable model of advanced brain ageing. Overall, it seems that structural ‘brain age’ in dogs, as in humans, better predicts neurological disease risk than chronological age.

## Introduction

The global burden of neurological disease has dramatically increased in the last 25 years, largely due to an ageing human population — a trend mirrored in companion animals.^1–2^ Much overlap exists between humans and domestic dogs with respect to age-linked vascular, degenerative, and neoplastic brain disorders. Shared environmental influences between these species, as well as the shorter lifespan and refined genetic architecture of pedigree dogs, has driven canines to the leading edge of comparative neurological disease modeling.^3–6^

Brain ageing varies among humans, and biological (physiological) ‘brain age’ better predicts disease risk than chronological age.^7–13^ These divergent ageing trajectories might be accentuated in the domestic dog, where selective breeding has produced extreme phenotypic diversity among pedigrees, and where longevity and the onset of age-related brain pathology is breed-dependent.^14–18^ Emerging evidence points to an increased risk of disease and mortality in humans with structurally ‘older’-appearing brains — dementia, epilepsy, and schizophrenia have all been associated with this enhanced ‘brain age’.^8–10,12–13,19–22^ Robust biomarkers of brain ageing are therefore of urgent clinical interest to identify individuals that deviate from a healthy ageing trajectory, enabling targeted early intervention.^8–10^

Certain brain morphometric parameters are predicted to change with neural decline.^8–9,16,18,23–41^ However, age-related structural changes are subtle, non-linear, and non-uniform in their distribution.^8,10,42–43^ Whilst a single measure is clinically convenient, it is unlikely to capture a phenotype for the complex biological process of ageing.^8–10^ Machine learning techniques that estimate brain age from human magnetic resonance imaging (MRI) data rely on the fact that morphometric correlates of brain ageing vary little between healthy individuals.^8^ This cannot be presumed in the dog, where breed morphometric variations present computational and statistical challenges.^5,14,44–45^ Isolating allometric (size-dependent) and non-allometric shape variation is problematic,^46–47^ and whilst automated MRI atlas-based protocols have emerged to assess canine brain morphometry,^44–45^ their accuracy remains inferior to manual morphometric extraction for dogs with different craniofacial morphologies.^5,29,44^ These morphologies — brachycephalic, mesocephalic, and dolichocephalic (‘short-headed, medium-headed, and long-headed’, respectively) — can impact as much on brain shape, as they do on external features of the head.^14,44^

Recent studies have addressed the phenotypic diversity of the domestic dog,^15,44–45,48–53^ but the morphometric diversity of the canine brain in a clinical context remains unexplored. Clinical datasets offer several advantages, not least that the natural progression of disease can be observed on a background of both individual- and breed-based heterogeneity. An obvious challenge in exploiting such data is its complexity. To address this issue, we have employed correlation-based network analysis, an unbiased, data-driven method used originally for analysis of transcriptomics data,^54–56^ and more recently to explore patient parameters associated with complex syndromes.^57^ A key attraction of network analysis is that it incorporates interactions within and between traits — as shown for behavioural phenotypes in dogs.^58^ Moreover, network analysis can test previous assumptions made about disease mechanisms and the clinical significance of patient-derived observations.^57,59^

In this study, we have applied network correlation analysis to a complex canine neurological dataset to explore how MRI-based brain morphometry profiles vary according to patient demographics and diagnosis. Our primary objective was to test statistically for co-enrichment between patient factors, clinical data, and brain morphometric features to extract novel insights into neurological disease risk.

## Materials and Methods

### Experimental design

The study objective was to conduct a large-scale, retrospective, unbiased analysis of canine brain morphometric data to derive features that might predict neurological disease risk in dogs. The Royal (Dick) School of Veterinary Studies Hospital for Small Animals data management system was screened for canine brain MRI scans performed between July 2009 and March 2017. The start date was dictated by MRI availability, and the end point when a minimum of 300 brain scans had been scheduled. Inclusion criteria were MRI of the whole brain, with at least one transverse and one sagittal sequence (T1-weighted or T2-weighted), and accessible clinical history. Patients with any trauma or procedure that would alter skull or brain morphometry were excluded. MRI scans were anonymized prior to blinded, quantitative data collection by one of two independent observers (observer A, O.M.S. and observer B, N.M.R.) using the same scoring protocol (Supplementary Figure S4). Analysis of 47 prospective scans (that met inclusion criteria) were used to assist with craniofacial category assignment by CFR, scored by observer B.

### Animals and ethics statement

MRI data were acquired from canine patients as part of routine diagnostic work-up. All patients had been referred to the Hospital and were assessed under the supervision of Board-certified specialists in Small Animal Internal Medicine and/or Neurology. Dogs were anaesthetized and scanned under the supervision of Board-certified specialists in Anaesthesia and Diagnostic Imaging, respectively. Written informed consent of each dog owner was obtained for all diagnostic procedures and for the use of anonymized clinical and imaging data for research purposes.

### Data acquisition

Body weight (kg) was extracted from the anaesthetic record on the day of MRI acquisition. Age was calculated using the date of birth and date of MRI acquisition. Meta-data (sex, breed, category of neurological diagnosis) were extracted using the clinical history, MRI report, clinical pathology reports, final neurologist report and (where available) histopathology reports. ‘Breed group’ categories were assigned according to the UK Kennel Club registration system (http://www.thekennelclub.org.uk); mixed breed dogs, and those without official breed recognition were either designated a ‘Crossbreed’ grouping or grouped according to the main contributing breed (e.g. Patterdale Terrier = Terrier; Collie X = Pastoral; Beagle X, Whippet X = Hound). Anomalous conditions included Chiari-like malformation, syringomyelia, hydrocephalus; inflammatory conditions were immune-mediated or infectious. The few dogs with degenerative myelopathy and normal brain MRIs were assigned a degenerative diagnosis. A ‘normal’ diagnostic category was assigned only in dogs with structurally normal brains where no neurological diagnosis was made (this contrasts with the study of Milne *et al*., where canine patients used for development of a brain atlas included dogs with ataxia, vestibular disease, and idiopathic cerebellitis).^44^ Brain morphometric features were assessed using OsiriX Medical Imaging Software, including previously published parameters and recognized normalization factors (Supplementary Figure S4). CFRs were derived using a modified version of the method described by Packer *et al*^60^ in which muzzle length (non-linear distance from dorsal tip of nasal planum to the stop in mm) is divided by cranial length (non-linear distance from occipital protuberance to the stop in mm). Measurements and precise locations of the nasal planum, stop and occipital protuberance were determined on mid-sagittal T2w images using the ‘open polygon’ tool of OsiriX and excluded obvious skin folds. Craniofacial categories were assigned based on (i) CFR (where available) and the cut-offs for craniofacial category assignment within our cohort or (ii) the average CFR available for that breed within our cohort. Brachycephaly was defined as a CFR of ≤ 0·52, mesocephaly as > 0·52 to < 0·67, and dolichocephaly as ≥ 0·67. Overall, 139 scans were scored by observer A and 172 scans by observer B, with an overlap of 25 scans to evaluate reproducibility of the scoring technique. Scoring between observers was highly reproducible (variance < 10%) for 8 parameters; for the remainder, scans measured by observer A were re-measured blind by the more experienced observer B (a board-eligible veterinary neurologist), before processing of the dataset for network analysis (Supplementary Figure S4e). For scans scored by both observers, only data extracted by observer B were used for subsequent analysis.

### Data processing

Raw data were processed prior to further analysis; brain length, cerebellar volume,^61^ cerebellar diameter, interthalamic adhesion height, corpus callosum thickness, and ventricular parameters were normalized to total brain volume (which included ventricular volume).^62^ Cranial length, brain width, total brain volume, and sulcus depth were normalized to body weight to control for allometric scaling.^62^ Cerebellar compression length, cerebellar compression index and obex position were normalized to head angle to control for patient positioning (Supplementary Figure S4). Corpus callosum angle was not normalized. Normalized total brain volumes were retained within the dataset for network analysis but head angle was excluded. Measured ventricle height created a markedly skewed data set due to the recorded ‘zero’ value in most patients. These measurements were categorized to indicate visual integrity of the septum pellucidum: 0 mm = ‘intact’, > 0 mm < 3mm = ‘minor’ loss, > 3 mm < 6mm = ‘moderate’ loss, > 6 mm < 10 mm = ‘severe’ loss, > 10mm = ‘absent’. ‘Septal integrity’ thus became an additional meta-data parameter. Age at MRI was categorized as follows: > 0 < 2 years = ‘Immature’, ≥ 2 < 4 years = ‘Young adult’, ≥ 4 < 8 years = ‘Middle-aged’, ≥ 8 < 10 years = ‘Mature’, ≥ 10 years = ‘Geriatric’. Magnitude of variance differed greatly between morphometric measurements, with the potential to disproportionately bias clustering of dogs according to the impact of one or a few parameters. To ensure fair representation of all parameters within the correlation analysis, all numerical data were median-centered for each parameter.

### Network analysis

Normalized, scaled and categorized data were imported into Graphia Professional (Kajeka Ltd., Edinburgh UK), a network analysis software package that calculates data matrices, supports graphical clustering, performs enrichment analyses and identifies patterns in large, complex datasets. The software was originally developed for the analysis of gene expression data, in which the correlation coefficient serves as a measure of co-expression between gene profiles and is used to define edges in a correlation network.^54^ In this case, a Pearson correlation was chosen to measure similarity between individual MRI scans based on normalized global brain morphometry measurements. The network graph created from the data (Supplementary Data Files S1-S3) was based on a user-defined correlation threshold of *r* = 0·7. This threshold was chosen to incorporate the maximum number of nodes (patient scans) with a minimum number of edges (correlations between patient scans). The measurements of thirteen animals in the cohort shared no correlation with other animals above this threshold and were absent from the graph. Network topology was determined by the number of correlations > *r* = 0·7 between all scans. The MCL clustering algorithm^57^ was used to subdivide the graph into discrete clusters of canine MRI scans sharing similar brain morphometric features. Granularity of the clustering (cluster size) is determined by the inflation value (MCLi). For this study, MCLi was set at 2·2 (smallest cluster size of three nodes). A detailed description and validation of the MCL algorithm can be found elsewhere (http://micans.org/mcl).^63^

### Enrichment and statistical analysis

Graphia Professional’s enrichment analysis uses Fisher’s exact test to determine the probability of a cluster’s composition occurring purely by chance, and offers tools to statistically confirm enrichment of a particular class. Since the canine brain data contained several classes for each MRI scan, Fisher’s exact was used to test each cluster for a disproportionately high representation of each class descriptor. Enrichment outputs include a heatmap and table providing the observed and expected number of members of each class descriptor within each cluster. The corresponding adjusted Fisher’s *P*-value represents how statistically unlikely it is for a class descriptor to occur within a cluster; the lower this value, the more significant the result, and the more brightly it is displayed on the heatmap. All other analyses were conducted in GraphPad Prism 7·0; one-way analysis of variance (ANOVA) was applied to determine differences between group means, and two-tailed t-tests were used for subsidiary comparisons between datasets of equal and unequal variance (determined by F-test). Linear regressions tested for significance between lines of best fit as described in figure legends, and Fisher’s exact test was used to assess odds ratios. In all dot plots, thick horizontal bars represent the median value, and asterisks refer to significant differences by t-test as indicated (\*\*\*\**P* < 0·0005, \*\*\**P* < 0·001, \*\**P* < 0·01, \**P* < 0·05).

### Data sharing

Anonymised DICOM files are available on request. Supplementary Data Files are deposited at Mendeley Data (doi:10.17632/y2f9272bbd.1). A trial version of Graphia Professional is free to download at https://kajeka.com/.

## Results

### Complexity within a canine referral cohort

A total of 9,438 morphometric and clinical data points were extracted from 286 MRI scans conducted on 281 individual dogs (Figure 1a, Supplementary Figure S1). These included 61 UK Kennel Club breeds and all seven recognized Kennel Club breed groups (Supplementary Figure S2, Supplementary Data File S1). The most common breeds in the cohort were Labrador Retriever (12·9%), Cavalier King Charles Spaniel (CKCS; 8·7%), and Boxer (6·3%); 52·7% of scans derived from male dogs and 65·8% of patients were neutered. Median age at MRI was 6·8 y (range 0·2-17·4 y) and median body weight was 18·1 kg (range 1·2-97·0 kg). The distribution of body weights and ages according to breed grouping highlighted the diversity within our cohort (Figure 1b, Supplementary Figure S3). Measurements to determine craniofacial ratios (CFRs)^60^ were possible in 117 retrospective scans, and in 17 prospective scans used to support craniofacial category assignment (Figure 1c, Supplementary Figures S2b, S4, Supplementary Data File S2). Based on defined cut-offs, 35·7% of MRI scans used for network analysis derived from brachycephalic, 50·7% from mesocephalic, and 13·6% from dolichocephalic dogs. Brachycephalic dogs had shorter brains relative to their cranial length and were predominantly found within Toy, Utility and Working groups (Figures 1d-f, Supplementary Figure S3d). Mesocephalic and dolichocephalic dogs were mainly found within Gundog and Hound groups, respectively. The distribution of MRI scans across craniofacial categories according to genetic clade^64^ (Supplementary Figure S5) identified a large contribution of European Mastiff, Retriever, and UK Rural clades to brachycephalic, mesocephalic, and dolichocephalic MRI scans, respectively. Overall, our cohort reflected the complex demographic of canines referred to neurology, on a background of current breed preferences among UK dog owners.

**Fig. 1.**
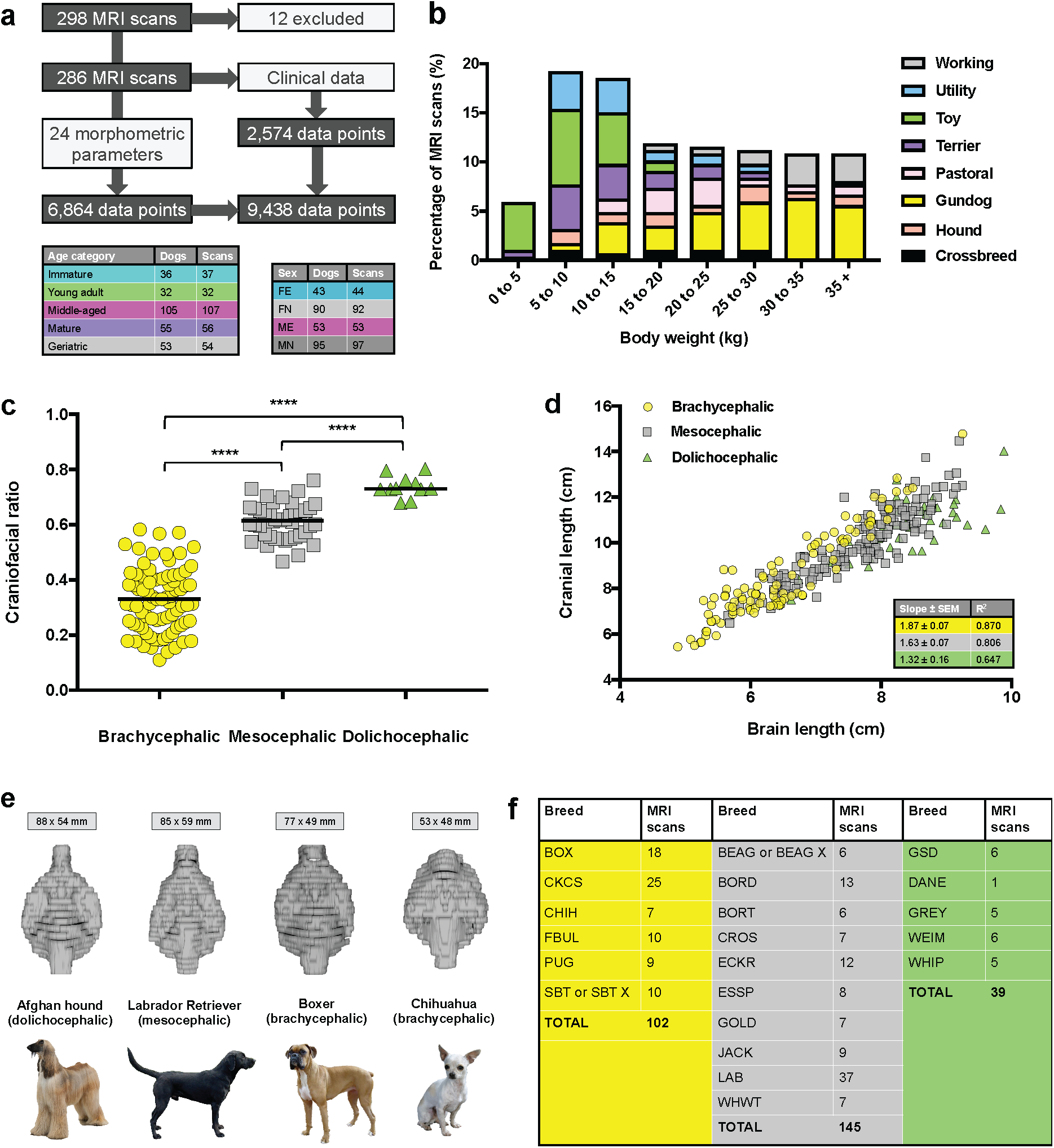
**A diverse and complex canine neurological cohort.** **(a)** Canine brain MRI scans; 12 were excluded due to lack of required MRI sequences; of the remaining 286 scans, 255 were from pure-bred dogs. Numbers of individual patients and MRI scans included in network analysis are tabulated. Five patients underwent two scans on separate dates. **(b)** Percentage of MRI scans by body weight according to breed group. **(c)** Measurable CFRs (screened from 333 MRI scans). Difference between group means was significant (*N* = 134; *P* < 0·0001; one-way ANOVA; \*\*\*\**P* < 0·0005 by t-test). **(d)** Linear regression of cranial length versus brain length according to craniofacial category. Differences between slopes were significant (*P* = 0·01 brachycephalic versus mesocephalic, *P* = 0·0007 brachycephalic versus dolichocephalic, *P* = 0·04 mesocephalic versus dolichocephalic); SEM = standard error of the mean. **(e)** Computed brain volumes for representative dolichocephalic, mesocephalic, and both large and small brachycephalic breeds; mean brain lengths and widths are shown for each breed. **(f)** Numbers of MRI scans used in network analysis for breeds with ≥ five representatives, by craniofacial category (brachycephalic, yellow; mesocephalic, grey; dolichocephalic, green). BOX (Boxer), CHIH (Chihuahua), FBUL (French Bulldog), PUG (PugDog), SBT (Staffordshire Bull Terrier), BEAG (Beagle), BORD (Border Collie), BORT (Border Terrier), CROS (Crossbreed), ECKR (English Cocker Spaniel), ESSP (English Springer Spaniel), GOLD (Golden Retriever), JACK (Jack Russell Terrier), LAB (Labrador Retriever), WHWT (West Highland White Terrier), GSD (German Shepherd Dog), DANE (Great Dane), GREY (Greyhound), WEIM (Weimeraner), WHIP (Whippet).

### Network analysis reveals clustering of canine brains

At a correlation threshold of *r* = 0·7, a graph was generated incorporating 273 MRI scans (nodes) and 3,911 correlations (edges) (Figure 2a, Supplementary Figure S6, Supplementary Data File S3). The graph’s topology exhibited distinct cliques (areas of high connectivity) and upon Markov clustering (MCL) produced 12 clusters, incorporating 250 scans (Supplementary Figure S7, Supplementary Table S1). Patients within each cluster shared similar brain morphometric features, with 71·3% of scans residing in one of six large clusters (Figure 2b). Figures 2c-e compare the brain morphometry profiles of the three most common breeds within the network; CKCS dogs were distinguished by their ventricular parameters and cerebellar compression, and Boxer and Labrador brains diverged mainly on the basis of ventricular size. To evaluate the statistical significance of cluster composition, an enrichment analysis was performed for each cluster of data (Figure 2f). Cluster one was enriched for brachycephalic Working dogs (including 14 Boxers). Immature and Chihuahua dogs were over-represented in cluster two, whilst cluster three was enriched for mesocephalic Gundogs including Labradors. Cluster four featured mesocephalic Crossbreed dogs, and mesocephalic dogs were also over-represented in cluster five. Dolichocephalic and geriatric dogs were enriched in clusters eight and eleven, respectively. Overall, relative to other craniofacial categories, brachycephalic dogs had larger brain widths, enlarged ventricular parameters, and greater cerebellar compression (Figures 3a-b, Supplementary Figure S8). Conversely, dolichocephalic dogs had narrower brains with intermediate ventricular volumes, and mesocephalic dogs had small ventricular volumes. Variation was observed in brain morphometry profiles according to sex; un-neutered animals had larger brains relative to their body weight, although this was offset by an increased ventricular size and sulcus depth in males (Figures 3c-d). Neutered and un-neutered females had the largest and smallest ventricular volumes, respectively. Whole brain parameters (length, width and volume), ventricular size, sulcus depth, and corpus callosum thickness separated the youngest and oldest dogs (Figures 3e-f). In summary, signalment (breed, craniofacial category, sex, and age) appeared to drive the clustering of canine brains, with ventricular size and brain width being most impacted by these factors.

**Fig. 2.**
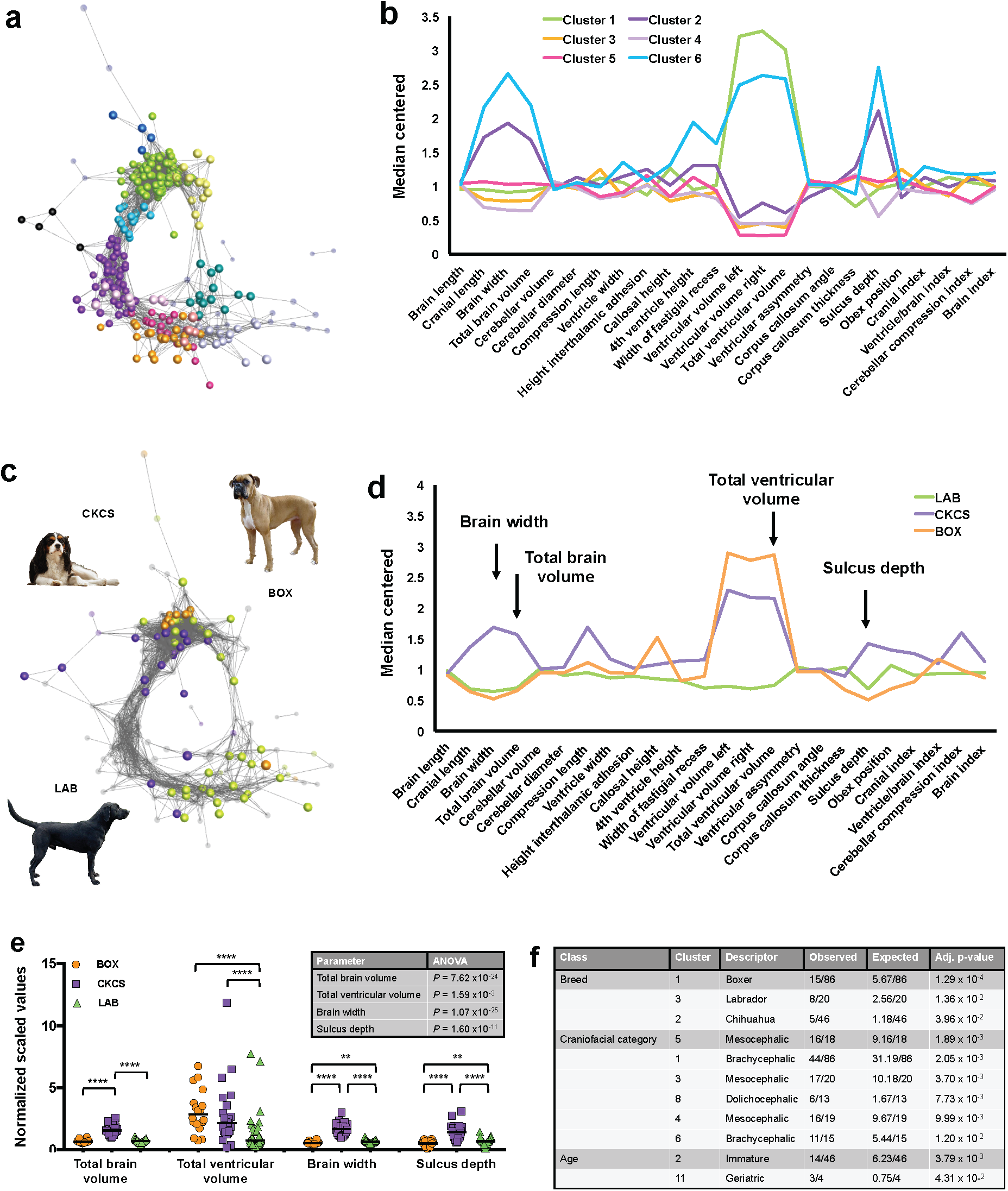
Network analysis reveals clustering of canine brains based on morphometry. In the network, nodes represent individual MRI scans; edges represent Pearson correlation coefficients (*r* > 0·7) between their brain morphometry profiles. Non-clustered and unselected nodes are displayed as smaller transparent spheres. Some nodes are hidden within clusters or on other aspects of the graph; iterations of the network can be explored by inputting Supplementary Data File S3 into Graphia Professional. **(a)** Network with nodes coloured by cluster; median lines for the six largest clusters are shown in associated chart **(b)**. Note that sulcus depth, ventricular volume, and whole brain parameters (length, width, volume) drive divergence of canine brain morphometry profiles. **(c-e)** Brain morphometry comparison for three most common breeds in the cohort. Arrows in **(d)** indicate key morphometric parameters tested in **E** (*N* = 79). Differences between group means were significant as shown in table (one-way ANOVA); in the dot plots, thick horizontal bars represent the median value, asterisks refer to significant differences by t-test (\*\*\*\**P* < 0·0005, \*\**P* < 0·01). **(f)** Enrichment analysis of breeds, craniofacial categories and age categories within clusters. Enrichments are listed only where observed node numbers were ≥ three (minimum cluster size). Note strong enrichment of brachycephalic Boxer dogs in cluster one.

**Fig. 3.**
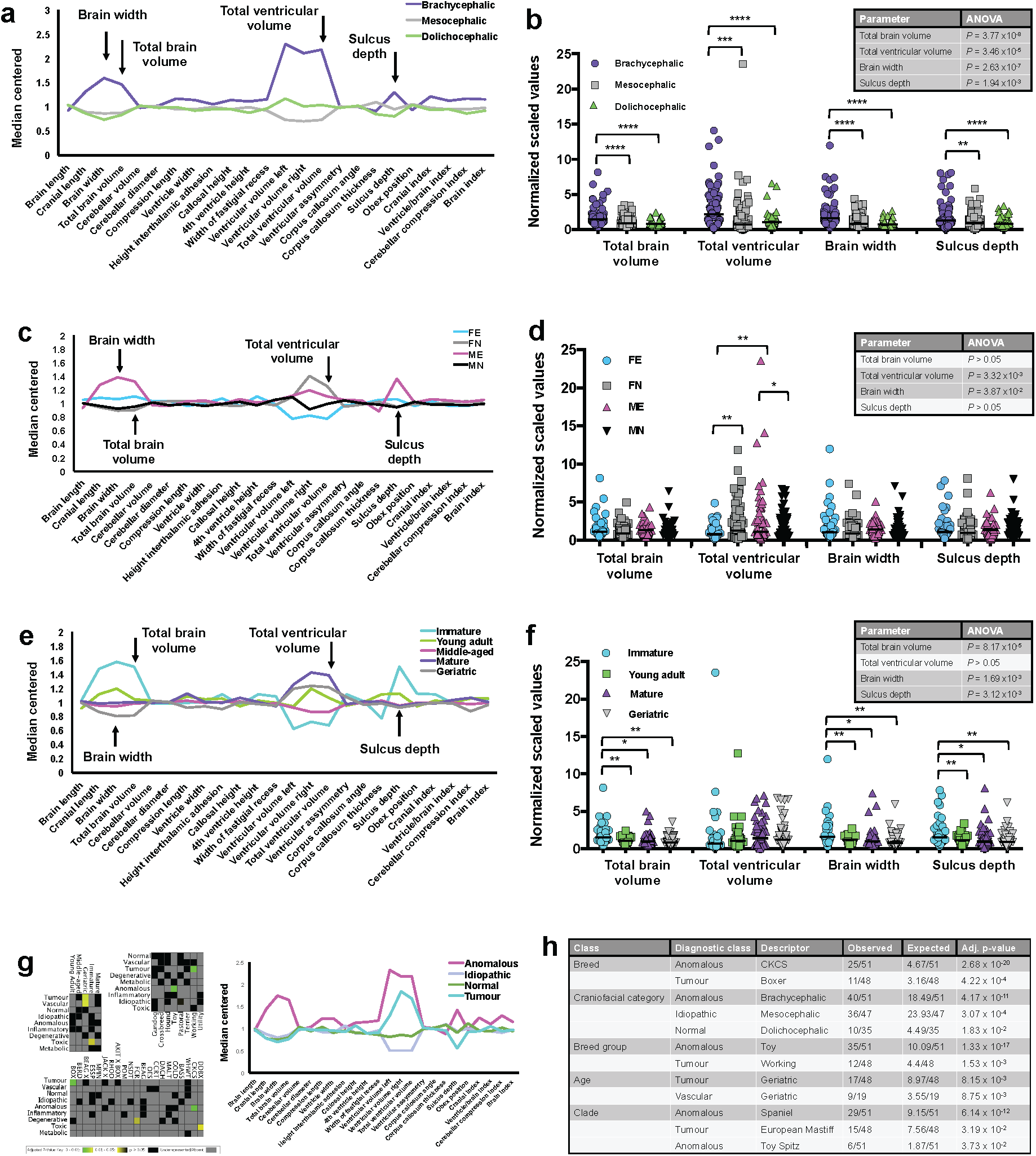
Signalment and diagnosis impact on canine brain morphometry. Brain morphometry comparisons by **(a-b)** craniofacial category, **(c-d)** sex, and **(e-f)** age category. Arrows in **a, c,** and **e** indicate key morphometric parameters tested in **b** (*N* = 286), **d** (*N* = 286), and **f** (*N* = 179) by one-way ANOVA. Differences between group means are shown in each inset table, with significance depicted in shaded boxes; asterisks refer to significant differences according to t-test (\*\*\*\**P* < 0·0005, \*\*\**P* < 0·001, \*\**P* < 0·01, \**P* < 0·05). In **c** and **d** FE = un-neutered females, FN = neutered females, ME = un-neutered males, and MN = neutered males. **(g)** Heat maps and chart coloured by final neurological diagnosis. Note enrichment of tumour diagnoses with both the geriatric group and Boxer breed. Network graphs for each diagnostic class are visualized separately in Supplementary Figure S10. **(h)** Enrichment analysis results for diagnostic class sets. Table lists significant enrichments together with expected and observed numbers for each descriptor that occurred in a given class, with adjusted *P*-values. Enrichments were excluded where observed number of nodes was < three (minimum cluster size). For each class, descriptors are listed in order of statistical significance. No enrichments were found within diagnostic class sets for septal integrity or sex.

### Clinical-morphometric interactions identify the Boxer as an outlier

Observing that some diagnostic classes were prominent among certain demographic categories and clusters (Figure 3g, Supplementary Figures S9-10), we next explored correlations between signalment, brain morphometry, and neurological disease. Interestingly, cluster one contained 26 dogs with tumour diagnoses (ten of which were Boxers) and there were patients in all breed groups with ‘idiopathic’ diagnoses based on clinical signs and a normal MRI — many of these had epilepsy. The Fisher’s exact test was used to detect enrichment of signalment descriptors within each diagnostic class (Figure 3h). Significant enrichments included brachycephalic dogs within the anomalous class, whilst geriatric dogs were enriched within tumour and vascular classes. Four out of ten Pointer Setter dogs had brain tumours, and three of these were neutered female geriatric Weimeraners (age and breed co-enriched with adjusted *P*-value of 2·38 x 10^-2^). Boxer dogs were greatly enriched within the tumour class, and mesocephalic dogs were over-represented within the idiopathic class. With respect to breed group, the anomalous class was significantly enriched for Toy dogs (mainly CKCS reflecting the high prevalence of Chiari-like malformation in this breed),^65^ whilst the tumour class was enriched for Working group dogs (mainly Boxers). Again, ventricular size strongly dictated clustering and group dynamics; four out of seven Labrador Retrievers positioned in cluster one had tumours and large ventricular parameters. Working and Toy breeds had the largest ventricular volumes, but these breed groups dramatically diverged with respect to whole brain parameters and sulcus depth. Strikingly, Boxers had remarkably narrow brains (Figure 4a), accentuating a feature more consistent with a dolichocephalic phenotype (Figures 3a). Moderate to severe loss of the septum pellucidum (membrane that separates the lateral ventricles of the brain) was prominent in the European Mastiff clade, which was also enriched for entire male dogs (adjusted *P*-value 9·55 x 10^-3^). Septal integrity was most compromised in the Boxer; only five out of 18 dogs had a visually intact septum (Supplementary Figure S11). Combined with ventriculomegaly, the reduced whole brain dimensions in the Boxer resulted in a small residual brain tissue volume relative to body size, a feature which clearly separated the Boxer from other brachycephalic breeds (Figure 4b). Together, our results defined the Boxer as an outlier, displaying both brachycephalic and dolichocephalic morphometric features, alongside an increased tumour risk.

**Fig. 4.**
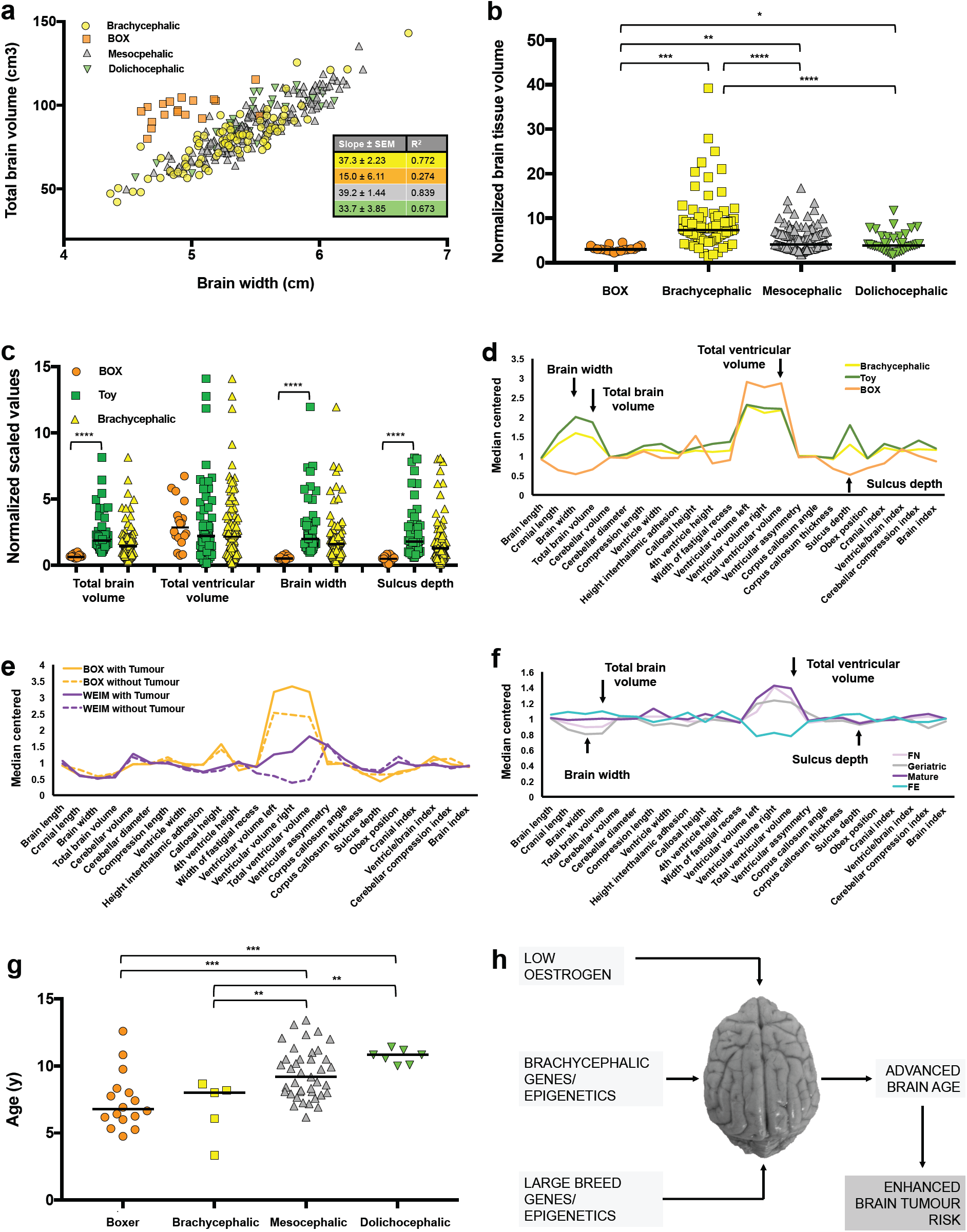
Advanced ‘brain age’ linked to tumour risk and oestrogen loss. **(a)** Linear regression of total brain volume versus brain width; differences between Boxer and other slopes were significant (*P* = 0·004 other brachycephalic, *P* = 0·0001 mesocephalic, *P* = 0·02 dolichocephalic); SEM = standard error of the mean. **(b)** Residual brain tissue volume (ventricular volume subtracted from total brain volume) normalized to body weight. Difference between group means was significant (*N* = 286, *P* < 0·0001; one-way ANOVA). **(c-d)** Total brain volume, brain width, and sulcus depth (arrowed) are small in the Boxer relative to other brachycephalic breeds. **(e)** The Boxer brain morphometry profile is not explained by tumour growth, which in another large breed (Weimeraner) has a marked impact on ventricular size and network position (Supplementary Figure S13G). **(f)** Age distribution of patients diagnosed with brain tumours since the start of the study period by craniofacial category. Difference between group means was significant (*N* = 67, *P* < 0·0001; one-way ANOVA). **(g)** Brain morphometry of neutered female dogs mimics that of mature and geriatric dogs, with a significantly larger ventricular volume than in un-neutered females. **(h)** Proposed model for factors contributing to advanced brain age and brain tumour risk in dogs.

### Advanced ‘brain age’ in the Boxer and neutered female dogs

Having confirmed enrichment of the Boxer breed with tumours, but not with the geriatric class (despite geriatric scans being enriched for tumours), we considered that Boxer brains may be subject to accelerated ageing. Indeed, Boxer brain morphometry profiles exaggerated those of mature and geriatric dogs (Figure 3e). Apart from ventriculomegaly, the ‘aged’ Boxer profile did not broadly represent the brachycephalic phenotype (Figure 4c-d). Boxer brain morphometric features were shared with some other members of the Working group (Rottweiler and Dogue de Bordeaux), and European Mastiff clade (Boston Terrier and Rhodesian Ridgeback; Supplementary Figure S12), but not all representatives (French Bulldog and Staffordshire Bull Terrier). To confirm that Boxer brain morphometry did not simply reflect tumour growth, the profiles of breeds with a high risk of tumours in our cohort (Boxer and Weimeraner) were compared, in the presence and absence of tumour diagnoses (Figure 4e). In the Boxer, tumour diagnosis was associated with a marginal increase in ventricular size, whereas in the Weimeraner, it converted a small ventricular profile to one consistent with ventriculomegaly. Follow-up scans in five dogs exposed the dynamism of cerebrospinal fluid (CSF)-filled spaces in response to partial or complete resolution of brain lesions (Supplementary Figure S13). However, the ‘aged’ morphometry profile appeared unique to the Boxer and was retained both before and after treatment. Intriguingly, the aged Boxer profile mimicked that of neutered females (Figure 4f), which had a high proportion of tumour diagnoses (21·1%) relative to un-neutered females (4·6%; the lowest percentage of the four sex categories within the network). By contrast, un-neutered females had relatively large whole brain parameters, small ventricles, and enriched with immature brain profiles (*P* = 8·97 × 10^-4^; Supplementary Figure S14). Critically, although neutered females were on average older than un-neutered females in our cohort, the relative increase in the size of their ventricles was significant in the geriatric group (Supplementary Figure S15). Within the network, seven of ten Boxer dogs with tumours were middle-aged. Prospective analysis of 148 dogs presenting for brain MRI at our institution identified an additional 23 dogs with brain tumours, all of which were entire males or neutered animals, including four Boxers with a mean age of 7·5 y. Boxers and other brachycephalic dogs were thus diagnosed earlier with brain tumours than other breed types (*P* < 0·01, Figure 4g). Finally, considering all brain scans performed to date (441 in 429 dogs), and excluding tumours that had metastasized from other parts of the body to the brain (Supplementary Data File S4), neutering increased the relative risk of brain tumours 11-fold in females (odds ratio 13·5, *P* = 0·0006, 95% confidence interval 2·4-141·4), and un-neutered females were seven times less likely to suffer brain tumours than un-neutered males (odds ratio 7·5, *P* = 0·03, 95% confidence interval 1·3-82·4). In conclusion, our findings suggest that oestrogen may be protective against brain ageing and brain tumour growth in dogs, whereas the Boxer is at high risk for both (Figure 4h).

## Discussion

Applying a data-driven approach, we have identified an aged-brain morphometric phenotype in Boxers and neutered female dogs that enriches with brain tumour risk. To our knowledge, this is the first network analysis of global brain morphometry, and the first study to link brain tumour risk to accelerated brain ageing. Our results are consistent with the hypothesis that structural brain age influences disease risk, and that oestrogen plays a role in brain ageing and tumour growth.

All canine patients underwent MRI because of brain-localising signs, therefore our cohort does not represent healthy brain ageing. Combining idiopathic and normal (undiagnosed) patients, our network included 31·3% structurally ‘normal’ brains. Subtle morphometric changes (Supplementary Figure S16) caution against describing idiopathic epileptic brains as ‘normal’,^66^ although others have used epileptic patients to establish reference values in dogs.^34^ Some tumour diagnoses were not confirmed by necropsy (Supplementary Data File S4), although where suspected, major differentials were ruled out with CSF analysis. Manual planimetry techniques are arguably more precise than semi-automated approaches,^42,44^ however the laborious measurement protocol used for this study is not appropriate for routine clinical application. Whilst templates derived from a small number of breeds without structural brain pathology enable rapid and reproducible morphometric analysis,^29,44–45,67^ these templates cannot accommodate the diverse brain morphologies observed in canine patients. Finally, referral bias will have magnified enrichments for breeds that most frequently present to our institution; network analysis partly controls for this issue, but it cannot eliminate the need for larger datasets to model disease risk at the population level.

A 40-fold difference in skeletal size exists between the largest and smallest dog breeds, and there is a strong correlation between body weight and the volume of several brain compartments.^34,50,62,68–69^ Most parameters were normalized to total brain volume to help control for individual and breed variations in brain morphometry.^25,30,70^ Some have argued against using ventricular-to-cerebrum ratio to assess brain ageing since such measures would be breed-specific,^62^ yet this presumes that ventriculomegaly reflects normal breed variation. Boxers without cerebral disease have large lateral ventricles relative to other breeds,^71^ however the definition, development, and clinical significance of ventriculomegaly in dogs remains controversial.^72^ Ageing has been associated with changes in brain and ventricular volume in dogs, but most data comes from laboratory Beagles.^16,18,23–24,26–27,29–30,35,73^ Given the extensive breed variation in canine ventricular morphology, age- and breed-specific reference ranges using a standard set of MRI sequences in neurologically normal dogs are needed. Training healthy dogs to participate in advanced neuroimaging studies without anaesthesia may address some ethical concerns and deliver the statistical power required for complex morphometric research questions.^5,42^

Certain observations built confidence in our analysis, not least the relative ventriculomegaly in brachycephalic breeds, and cerebellar compression in CKCS dogs.^65,72,74^ Enrichment of Boxers with tumours was anticipated^74^ and the compromised septal integrity in this breed is more common in brachycephalics generally (29% versus 9% and 13% in mesocephalic and dolichocephalic breeds in our network, respectively). Non-detection of the canine septum pellucidum on MRI is largely considered incidental,^70,75–76^ and it remains possible that the septum is intact, but too thin to be observed in some dogs.^70^ Apparent absence of the septum has been observed in neurologically normal humans but is often associated with other structural anomalies.^70,77^ Interestingly, 21 of 25 CKCS dogs in our cohort had intact septa, despite their high prevalence of Chiari-like malformation.^78^ Conceptually, a compromised septum might increase ventricular compliance and thus explain why Boxers are at low risk of Chiari-like malformation and syringomyelia, despite shared ventricular morphology with CKCSs.

The need to explore sexual dimorphism in brain ageing is underpinned by the fact that dementia disproportionately affects women.^79^ The largest ever single-sample neuroanatomical study of sex differences using UK Biobank data found several sexually dimorphic differences in human brain structure.^42^ Importantly these changes operated in a global manner, supporting our approach to consider multiple morphometric features in concert, and to correct for total brain volume. Age-related structural brain changes differ between men and women,^36^ and also between male and female dogs.^16,29^ Men exhibit greater increases in sulcal and ventricular CSF volume,^36^-^37^ whilst women demonstrate greater rates of hippocampal atrophy.^38–40^ A semi-quantitative visual rating scale was used to chart cerebral involutional changes in dogs, however neither sex nor neutering status were considered as co-variates.^23^ One canine study reported that different brain regions appeared more vulnerable to atrophy in males — although these animals were all sexually intact.^29^ Post-mortem studies in German Shepherd Dogs found ventricular enlargement with ageing and no apparent relationship to sex, although again the effect of neutering was not explored.^27^ By contrast, our results indicate an accelerated ventriculomegaly and total brain loss in neutered female dogs. Importantly, whilst there was a difference in age distribution between sex categories in our network, there was a trend for enhanced ventriculomegaly in neutered females across all age categories in adulthood, reaching significance in the geriatric group. To the authors’ knowledge, this is the first study to demonstrate an effect of neutering status on brain morphometry in dogs.

Oestrogen deficiency is proposed to explain accelerated brain ageing in post-menopausal women as well as an accelerated epigenetic clock in ovariectomized mice.^80–82^ Several meta-analyses have shown hormone-replacement therapy (HRT) to be neuroprotective, and although recent publications have raised doubt over the ‘oestrogen deficiency’ theory of dementia,^83–85^ HRT may still defend cognition in a subgroup of women in the perimenopausal period.^85–86^ Our work supports the concept that oestrogen loss may accelerate neural decline, however causal mechanistic insight is lacking. Development of multicentre canine biobanks will facilitate investigations of oestrogen status as a function of brain ageing in dogs, and whether this relates to cognitive dysfunction. In human patients, advanced structural ‘brain-age’ is often paralleled by epigenetic markers of ageing.^9,11,87–88^ The premise that oestrogen may have neuroprotective benefits across the lifespan — and that its effects may be epigenetically regulated^89^ — emphasises the need to integrate structural, functional, and molecular approaches in the study of brain ageing and brain disease.

Canine gliomas occur most commonly in brachycephalic breeds, with the Boxer at highest risk.^74^ We noted a significant enrichment of brain tumours with Boxers, however the absence of tumour diagnoses in CKCS dogs resulted in non-enrichment of tumours within our brachycephalic category. The majority of Boxers in our cohort were middle-aged, consistent with Song *et al* where gliomas most frequently occurred in dogs aged seven-to-eight years.^74^ This is despite the fact that increasing age remained a risk factor for all intracranial neoplasias (as seen here).^74^ An increased risk of primary intracranial neoplasms has also been found in large breed dogs,^74^ and indeed, only 11 of 52 dogs with tumour diagnoses in our network were small breeds. Most size variation between purebred dogs is controlled by a few genes of major effect, including several members of the insulin-like growth factor-1 (*Igf-1*) pathway.^4,46,50^ *Igf-1* is a major determinant of dog size; its variable expression is proposed to underlie the increased longevity of smaller breeds and the higher frequency of neoplasia-associated deaths in large breeds.^62^ Coincidentally, the rapid growth of large breeds may initiate premature ageing due to increased free radical release during development.^90^ Roughly half of small or medium breed dogs also have ‘large alleles’, mainly found in muscled breeds such as the French Bulldog and Boxer.^46,48^ Our analysis reveals that Boxers have ventricular parameters of the small brachycephalic phenotype, but whole brain parameters of large breed mesocephalic or dolichocephalic phenotypes. Conceivably, a combination of variants promoting brachycephaly (e.g. *Smoc2*), on a background of those promoting growth (*Igf-1*) may place the Boxer at extreme risk of premature ageing and brain tumours.^48^

Primary brain tumours, including malignant gliomas, are more common in men globally, indicating that sex plays a role in brain tumour pathogenesis.^91^-^92^ Furthermore, the risk of intracranial tumours is increased in women with complete or partial X-chromosome monosomy and low oestrogen levels.^93^ The human male predominance for brain tumours persists in all age groups, indicating that acute effects of circulating sex steroids cannot simply explain the sexual disparity in tumour risk.^91^ Mosaic loss of chromosome Y, the most common acquired human mutation and another putative biomarker of ageing, has been associated with an increased risk both of Alzheimer’s disease and various cancers.^94–96^ Emerging evidence thus supports a role for both chromosomal and gonadal sex in neuro-oncogenesis and brain ageing.^97^ To our knowledge, this is the first study to report a reduced risk of brain tumours in un-neutered female dogs relative to neutered animals. Given the routine (but not mandatory) practice of neutering, an unrivalled opportunity exists to explore the influence of chromosomal and gonadal sex on neuropathology in canines of varied neutering status.

Extreme breed characteristics impact on health and welfare, with widespread concerns surrounding brachycephaly.^61,90^ Our work extends this to the brain, highlighting an urgency to better understand the factors that influence brain ageing in dogs. Simultaneously, comparative studies will accelerate our knowledge of how chromosomal and hormonal sex affect brain structure, brain ageing, and brain tumour development in humans. The Boxer breed in particular could represent a valuable model of naturally-enhanced brain ageing. Larger, longitudinal imaging studies are required to confirm how patient demographics influence brain age — network analysis can facilitate discovery of subtle yet important phenotypic shifts within these complex clinical datasets. Importantly, our unique application of network analysis can be immediately translated to pre-existing and emerging human patient data. A key question is whether canine brain morphometry and associated morbidity can be explained by selectively-driven changes in skull shape, or whether independent genetic, epigenetic, or epidemiological factors contribute to neurological disease. Isolating these factors will advance our understanding of disease pathogenesis, with important implications for canine and human brain health.^3,6^

## Acknowledgments

The authors gratefully acknowledge the Diagnostic Imaging Service and Dr Darren Shaw of the Royal (Dick) School of Veterinary Studies for technical and advisory support. MRI scanning facilities were provided by Burgess Diagnostics Ltd., Preston, UK. Graphia Professional is a software product of Kajeka Ltd., Edinburgh, UK. OsiriX Medical Imaging Software was developed by Pixmeo SARL, Bernex, Switzerland. Images of canine breeds were all downloaded via Wikimedia Commons and each modified to remove background image data. Attributions are as follows: AFGH (By SheltieBoy (Flickr: AKC Helena Fall Dog Show 2011) [CC BY 2.0 (https://creativecommons.org/licenses/by/2.0)]); BOX (By Flickr user boxercab (Flickr here) [CC BY 2.0 (https://creativecommons.org/licenses/by/2.0)]); CHIH (Photo taken by en:User:Exdumpling in 2004 and uploaded to English Wikipedia as WhiteTanChihuahua.jpg claiming own work with PD-self license); FBUL (By The original uploader was EGILEO at Italian Wikipedia [CC BY 2.0 (https://creativecommons.org/licenses/by/2.0)]; DDBX (By X posid [CC0]); GREY (By FLickr user Scott Feldstein (Flickr here) [CC BY 2.0 (https://creativecommons.org/licenses/by/2.0)]); LAB (By Desaix83, d’après le travail de Chrizwheatley [Public domain]); Shiba Inu (By Takashiba at English Wikipedia [Public domain]); SPOO (By Inbalsigal [CC BY 3.0 (https://creativecommons.org/licenses/by/3.0)]).

## Role of funding source

This work was supported by a Wellcome Trust Integrated Training Fellowship for Veterinarians (096409/Z/11/Z to N.M.R) and an MSD Animal Health Connect Bursary (to O.M.S.). Funding sources did not have any involvement in the study design; the collection, analysis and interpretation of data; writing of the report; or the decision to submit the article for publication. The corresponding author had full access to all of the data used in the study and is responsible for the decision to submit this work for publication.

## Declaration of interests

T.F. is a company director of Kajeka Ltd. registered at Sir Alexander Robertson Building, University of Edinburgh, Easter Bush Campus, Edinburgh, EH25 9RG, UK. The authors declare no competing interests.

## Author contributions

Conceptualization, N.M.R; Methodology, N.M.R., O.M.S., T.S., J.J.S. and T.F.; Data acquisition, N.M.R., O.M.S., and J.J.S.; Formal Analysis, N.M.R, O.M.S., J.J.S. and T.F.; Investigation and interpretation, N.M.R, O.M.S., L.H., K.M-H, D.J.A., J.J.S., and T.F.; Resources, N.M.R, K.M-H, T.S., D.J.A., J.J.S. and T.F.; Writing – original draft, N.M.R.; Writing – review and editing, N.M.R, O.M.S., L.H., K.M-H., T.S., D.J.A., J.J.S. and T.F.; Visualization, N.M.R., O.M.S., J.J.S. and T.F.; Supervision, N.M.R, and T.F.; Project administration, N.M.R.; Funding acquisition, N.M.R., O.M.S., and D.J.A. All authors approved the final version to be submitted.

## Supplementary Materials

### Supplementary Figures

Fig.S1. Patient parameters recorded for each MRI scan Fig.S2. Breed characteristics for MRI scans

Fig.S3. Age and body weight distribution of dogs in network

Fig.S4. Canine brain morphometric measurement protocol and reproducibility Fig.

S2. Percentage of MRI scans by genetic clade

Fig.S6. Correlation graph settings for Graphia Professional dataset Fig.

S7. Cluster characteristics

Fig.S8. Networks by signalment

Fig.S9. Demographics and enrichment of diagnoses Fig.S10. Diagnostic class morphometrics

Fig.S11. Septal integrity impacts on brain morphometry

Fig.S12. Shared morphometric features of the Boxer

Fig.S13. Patients scanned twice during the study period

Fig.S14. Enrichments by sex

Fig.S15. Ventricular enlargement in neutered female dogs

Fig.S16. Dichotomous brain morphometry in dogs with and without paroxysmal events

## Supplementary Tables

Table S1. Clusters generated for network

**Supplementary Data Files**

Data File S1. Raw data used for network analysis

Data File S2. CFRs used to support craniofacial category assignment Data File

S3. CSV. file for network analysis

Data File S4. All scans and tumour diagnoses

## References

1. GBD 2015 Neurological Disorders Collaborator Group. Global, regional, and national burden of neurological disorders during 1990-2015: a systematic analysis for the Global Burden of Disease Study 201Lancet Neurol. 2017; 16: 877–897.

2. Day MJ. Ageing, immunosenescence and inflammageing in the dog and cat . J. Comp. Pathol. 2010; 142: Suppl 1:S60–9.

3. Kol A, Arzi B, Athanasiou KA, et al. Companion animals: Translational scientist’s new best friends. Sci. Transl. Med. 2015; 7: 308ps21.

4. Hayward JJ, Castelhano MG, Oliveira KC, et al. Complex disease and phenotype mapping in the domestic dog. Nat. Commun. 2016; 7: 10460.

5. Bunford N, Andics A, Kis A, Miklósi Á, Gácsi M. Canis familiaris as a model for non-invasive comparative neuroscience. Trends Neurosci. 2017; 40: 438–452.

6. Christopher MM. One health, one literature: Weaving together veterinary and medical research. Sci. Transl. Med. 2015; 7: 303fs36.

7. Jylhävä J, Pedersen NL, Hägg S. Biological Age Predictors. EBioMedicine 2017; 21: 29–36.

8. Cole JH, Franke K. Predicting Age Using Neuroimaging: Innovative Brain Ageing Biomarkers. Trends Neurosci. 2017; 40: 681–690.

9. Cole JH, Ritchie SJ, Bastin ME, et al. Brain age predicts mortality. Mol. Psychiatry, 2018; 13: 1385–1392.

10. Cole JH, Marioni RE, Harris SE, Deary IJ. Brain age and other bodily ‘ages’: implications for neuropsychiatry. Mol. Psychiatry 2018; doi:10.1038/s41380-018- 0098-[Epub ahead of print].

11. Levine ME, Lu AT, Bennett DA, Horvath S. Epigenetic age of the pre-frontal cortex is associated with neuritic plaques, amyloid load, and Alzheimer’s disease related cognitive functioning. Aging (Albany NY), 2015; 7: 1198–1211.

12. Schnack HG, van Haren NE, Nieuwenhuis M, Hulshoff Pol HE, Cahn W, Kahn RS. Accelerated Brain Aging in Schizophrenia: A Longitudinal Pattern Recognition Study. Am. J. Psychiatry, 2016; 173: 607–616.

13. Koutsouleris N, Davatzikos C, Borgwardt S. Accelerated brain aging in schizophrenia and beyond: a neuroanatomical marker of psychiatric disorders. Schizophr. Bull. 2014; 40: 1140–1153.

14. Roberts T, McGreevy P, Valenzuela M. Human induced rotation and reorganization of the brain of domestic dogs. PLoS One 2010; 5: e11946.

15. Boyko AR, Quignon P, Li L, et al. A simple genetic architecture underlies morphological variation in dogs. PLoS Biol. 2010; 8: e1000451.

16. Youssef SF, Capucchio MT, Rofina JE, et al. Pathology of the Aging Brain in Domestic and Laboratory Animals, and Animal Models of Human Neurodegenerative Diseases. Vet Path. 2016; 53: 327–348.

17. Greer KA, Canterberry SC, Murphy KE. Statistical analysis regarding the effects of height and weight on life span of the domestic dog. Res. Vet. Sci. 2007; 82: 208–14.

18. Vite CH, Head E. Aging in the canine and feline brain. Vet Clin. North Am. Small Anim. Pract. 2014; 44: 1113–29.

19. Gaser C, Franke K, Klöppel S, Koutsouleris N, Sauer H. Alzheimer’s Disease Neuroimaging Initiative, BrainAGE in Mild Cognitive Impaired Patients: Predicting the Conversion to Alzheimer’s Disease. PLoS One. 2013; 8: e67346.

20. Habes M, Janowitz D, Erus G. Advanced brain aging: relationship with epidemiologic and genetic risk factors, and overlap with Alzheimer disease atrophy patterns. Transl. Psychiatry 2016; 6: e775.

21. Löwe LC, Gaser C, Franke F. Alzheimer’s Disease Neuroimaging Initiative, The Effect of the APOE Genotype on Individual BrainAGE in Normal Aging, Mild Cognitive Impairment, and Alzheimer’s Disease. PLoS One 2016; 11: e0157514.

22. Pardoe HR, Cole JH, Blackmon K, Thesen T, Kuzniecky R. Human Epilepsy Project Investigators, Structural brain changes in medically refractory focal epilepsy resemble premature brain aging. Epilepsy Res. 2017; 133: 28–32.

23. Pugliese M, Carrasco JL, Gomez-Anson B, et al. Magnetic resonance imaging of cerebral involutional changes in dogs as markers of aging: an innovative tool adapted from a human visual rating scale. Vet. J. 2009; 186: 166–171.

24. Su MY, Head E, Brooks WM. Magnetic resonance imaging of anatomic and vascular characteristics in a canine model of human aging. Neurobiol. Aging 1998; 19: 479–85.

25. Su MY, Tapp PD, Vu L, et al. A longitudinal study of brain morphometrics using serial magnetic resonance imaging analysis in a canine model of brain aging. Prog. Neuropsychopharmacol. Biol. Psychiatry. 2005; 29: 389–97.

26. Kimotsuki T, Nagaoka T, Yasuda M, Tamahara S, Matsuki N, Ono K. Changes of magnetic resonance imaging on the brain in beagle dogs with aging. J. Vet. Med. Sci. 2005; 67: 961–967.

27. Gonzalez-Soriano J, Marin Garcia P, Contreras-Rodriguez J, Martínez-Sainz P, Rodríguez-Veiga E. Age-related changes in the ventricular system of the dog brain. Ann. Anat. 2001; 183: 283–91.

28. Tapp PD, Siwak CT, Gao FQ, et al. Frontal lobe volume, function, and beta-amyloid pathology in a canine model of aging. J. Neurosci. 2004; 24, 8205–13.

29. Tapp PD, Head K, Head E, Milgram NW, Muggenburg BA, Su MY. Application of an automated voxel-based morphometry technique to assess regional gray and white matter brain atrophy in a canine model of aging. Neuroimage. 2006; 29: 234–44.

30. Hasegawa D, Yayoshi N, Fujita Y, Fujita M, Orima H. Measurement of interthalamic adhesion thickness as a criteria for brain atrophy in dogs with and without cognitive dysfunction (dementia). Vet. Radiol. Ultrasound 2005; 46: 452–457.

31. Noh D, Choi S, Choi H, Lee Y, Lee K. Evaluation of interthalamic adhesion size as an indicator of brain atrophy in dogs with and without cognitive dysfunction. Vet. Radiol. Ultrasound. 2017; 58: 581–587.

32. Pilegaard AM, Berendt M, Holst P, Møller A, McEvoy FJ. Effect of Skull Type on the Relative Size of Cerebral Cortex and Lateral Ventricles in Dogs. Front. Vet. Sci. 2017; 4: 30.

33. Henke D, Böttcher P, Doherr MG, Oechtering G, Flegel T. Computer-assisted magnetic resonance imaging brain morphometry in American Staffordshire Terriers with cerebellar cortical degeneration. J. Vet. Intern. Med. 2008; 22: 969–75.

34. Thames RA, Robertson ID, Flegel T. Development of a morphometric magnetic resonance image parameter suitable for distinguishing between normal dogs and dogs with cerebellar atrophy. Vet. Radiol. Ultrasound. 2010; 51: 246–53.

35. Borràs D, Ferrer I, Pumarola M. Age-related changes in the brain of the dog, Vet. Pathol. 1999; 36: 202–11.

36. Coffey CE, Lucke JF, Saxton JA, et al. Sex differences in brain aging: a quantitative magnetic resonance imaging study. Arch. Neurol. 1998; 55: 169–79.

37. Pfefferbaum A, Rohlfing T, Rosenbloom MJ, Chu W, Colrain IM, Sullivan EV. Variation in longitudinal trajectories of regional brain volumes of healthy men and women (ages 10 to 85 years) measured with atlas-based parcellation of MRI. Neuroimage. 2013; 65: 176–93.

38. Murphy DG, DeCarli C, McIntosh AR, et al. Sex differences in human brain morphometry and metabolism: an in vivo quantitative magnetic resonance imaging and positron emission tomography study on the effect of aging. Arch. Gen. Psychiatry. 1996; 53: 585–94.

39. Crivello F, Tzourio-Mazoyer N, Tzourio C, Mazoyer B. Longitudinal assessment of global and regional rate of grey matter atrophy in 1,172 healthy older adults: modulation by sex and age. PLoS One 2014; 9: e114478.

40. Goto M, Abe O, Miyati T, et al. 3 Tesla MRI detects accelerated hippocampal volume reduction in postmenopausal women. J. Magn. Reson. Imaging. 2011; 33: 48–53.

41. Resnick SM, Pham DL, Kraut MA, Zonderman AB, Davatzikos C. Longitudinal magnetic resonance imaging studies of older adults: a shrinking brain. J. Neurosci. 2003; 23: 3295–301.

42. Ritchie SJ, Cox SR, Shen X, et al. Sex Differences In The Adult Human Brain: Evidence From 5,216 UK Biobank Participants. Cerebral Cortex, 2018; 28: 2959–2975.

43. Walhovd KB, Westlye LT, Amlien I, et al. Consistent neuroanatomical age-related volume differences across multiple samples. Neurobiol. Aging, 2011; 32: 916–932.

44. Milne ME, Steward C, Firestone SM, Long SN, O’Brien TJ, Moffat BA. Development of representative magnetic resonance imaging-based atlases of the canine brain and evaluation of three methods for atlas-based segmentation. Am. J. Vet. Res. 2016; 77: 395–403.

45. Nitzsche B, Boltze J, Ludewig E, et al. A stereotaxic breed-averaged, symmetric T2w canine brain atlas including detailed morphological and volumetrical data sets. Neuroimage 2018; pii S1053–8119: 30066–doi:10.1016/j.neuroimage.2018.01.06[Epub ahead of print].

46. Schoenebeck JJ, Ostrander EA. The genetics of canine skull shape variation. Genetics 2013; 193: 317–25.

47. Reardon PK, Seidlitz J, Vandekar S, et al. Normative brain size variation and brain shape diversity in humans. Science, 2018; 360: 1222–1227.

48. Marchant TW, Johnson EJ, McTeir L, et al. Canine Brachycephaly Is Associated with a Retrotransposon-Mediated Missplicing of SMOCCurr. Biol. 2017; 27: 1573–1584.e6.

49. Rimbault M, Beale HC, Schoenebeck JJ, et al. Derived variants at six genes explain nearly half of size reduction in dog breeds. Genome Res. 2013; 23: 1985–95.

50. Plassais J, Rimbault M, Williams FJ, Davis BW, Schoenebeck JJ, Ostrander EA. Analysis of large versus small dogs reveals three genes on the canine X chromosome associated with body weight, muscling and back fat thickness. PLoS Genet. 2017; 13: e1006661.

51. Bannasch D, Young A, Myers J, et al. Localization of canine brachycephaly using an across breed mapping approach. PLoS One 2010; 5: e9632.

52. Schoenebeck JJ, Hutchinson SA, Byers A, et al. Variation of BMP3 contributes to dog breed skull diversity. PLoS Genet. 2012; 8: e1002849.

53. Schoenebeck JJ, Ostrander EA. Insights into morphology and disease from the dog genome project. Annu. Rev. Cell Dev. Biol. 2014; 30: 535–60.

54. Freeman TC, Goldovsky L, Brosch M, et al. Construction, visualisation, and clustering of transcription networks from microarray expression data. PLoS Comput Biol. 2007; 3: 2032–2042.

55. Theocharidis A, van Dongen S, Enright AJ, Freeman TC. Network visualization and analysis of gene expression data using BioLayout Express (3D). Nat Protoc. 2009; 4: 1535–50.

56. Hume DA, Summers KM, Raza S, Baillie JK, Freeman TC. Functional clustering and lineage markers: insights into cellular differentiation and gene function from large-scale microarray studies of purified primary cell populations. Genomics 2010; 95: 328–338.

57. Hall DP, MacCormick IJ, Phythian-Adams AT, et al. Network analysis reveals distinct clinical syndromes underlying acute mountain sickness. PLoS One 2014; 9: e81229.

58. Goold C, Vas J, Olsen C, Newberry RC. Using network analysis to study behavioural phenotypes: an example using domestic dogs. R. Soc. Open Sci. 2016; 3: 160268.

59. Baillie JK, Bretherick A, Haley CS. Shared activity patterns arising at genetic susceptibility loci reveal underlying genomic and cellular architecture of human disease. PLoS Comput Biol. 2018; 14: e1005934.

60. Packer RM, Hendricks A, Tivers MS, Burn CC. Impact of Facial Conformation on Canine Health: Brachycephalic Obstructive Airway Syndrome. PLoS One 2015; 10: e0137496.

61. Driver CJ, Chandler K, Walmsley G, Shihab N, Volk HA. The association between Chiari-like malformation, ventriculomegaly and seizures in Cavalier King Charles Spaniels. Vet. J. 2013; 195: 235–7.

62. Schmidt MJ, Amort KH, Failing K, Klingler M, Kramer M, Ondreka N. Comparison of the endocranial- and brain volumes in brachycephalic dogs, mesaticephalic dogs and Cavalier King Charles Spaniels in relation to their body weight. Acta Vet. Scand. 2014; 56: 30.

63. van Dongen S, Graph clustering by flow simulation, PhD Thesis, University of Utrecht 2000.

64. Parker HG, Dreger DL, Rimbault M, et al. Genomic Analyses Reveal the Influence of Geographic Origin, Migration, and Hybridization on Modern Dog Breed Development. Cell Rep. 2017; 19: 697708.

65. Knowler SP, Cross C, Griffiths S. Use of Morphometric Mapping to Characterise Symptomatic Chiari-Like Malformation, Secondary Syringomyelia and Associated Brachycephaly in the Cavalier King Charles Spaniel. PLoS One 2017; 12: e0170315.

66. Whelan CD, Altmann A, Botia JA, et al. Structural brain abnormalities in the common epilepsies assessed in a worldwide ENIGMA study. Brain 2018; 141: 391–408.

67. Frank L, Lüpke M, Kostic D, Löscher W, Tipold A. Grey matter volume in healthy and epileptic beagles using voxel-based morphometry – a pilot study. BMC Vet. Res. 2018; 14: 50 (2018).

68. Reinitz LZ, Bajzik G, Garamvölgyi R, et al. Linear relationship found by magnetic resonance imaging between cerebrospinal fluid volume and body weight in dogs. Acta Vet. Hung. 2017; 65: 1–12.

69. Carreira LM. Using Bronson Equation to Accurately Predict the Dog Brain Weight Based on Body Weight Parameter. Vet. Sci. 2016; 3: pii E36.

70. Pivetta M, De Risio L, Newton R, Dennis R. Prevalence of lateral ventricle asymmetry in brain MRI studies of neurologically normal dogs and dogs with idiopathic epilepsy. Vet. Radiol. Ultrasound. 2013; 54: 516–21.

71. Schroder H, Meyer-Lindenberg A, Nolte I. Comparative examination of the lateral cerebral ventricles of different dog breeds using quantitative computed tomography. Berl. Munch Tierarztl. Wochenschr. 2006; 119: 506–511.

72. Schmidt MJ, Laubner S, Kolecka M, et al. Comparison of the Relationship between Cerebral White Matter and Grey Matter in Normal Dogs and Dogs with Lateral Ventricular Enlargement. PLoS One 2015; 10: e0124174.

73. Reifinger M. Volumetric examination of senile brain involution in dogs, Anat Histol. Embryol. 1997; 26: 141–6.

74. Song RB, Vite CH, Bradley CW, Cross JR. Postmortem evaluation of 435 cases of intracranial neoplasia in dogs and relationship of neoplasm with breed, age, and body weight. J. Vet. Intern. Med. 2013; 27: 1143–52.

75. Vite CH, Insko EK, Schotland HM, Panckeri K, Hendricks JC. Quanti?cation of cerebral ventricular volume in English bulldogs. Vet. Radiol. Ultrasound 1997; 38: 437–443.

76. De Haan CE, Kraft SL, Gavin PR, Wendling LR, Griebeno ML. Normal variation in size of the lateral ventricles of the Labrador Retriever dog as assesses by magnetic resonance imaging. Vet. Radiol. Ultrasound 1994; 35: 83–86.

77. Sundarakumar DK, Farley SA, Smith CM, Maravilla KR, Dighe MK, Nixon JN. Absent cavum septum pellucidum: a review with emphasis on associated commissural abnormalities. Pediatr. Radiol. 2015; 45: 950–64.

78. Driver CJ, Rusbridge C, Cross HR, McGonnell I, Volk HA. Relationship of brain parenchyma within the caudal cranial fossa and ventricle size to syringomyelia in cavalier King Charles spaniels. J. Small Anim. Pract. 2010; 51: 382–386.

79. Mazure CM, Swendsen J. Sex differences in Alzheimer’s disease and other dementias. Lancet Neurol. 2016; 15: 451–452.

80. Birge SJ. Hormones and the aging brain. Geriatrics 1998; 53: S28–30.

81. Melton L. Oestrogen shields brain from ageing. Lancet 1999; 354: 1101.

82. Stubbs TM, Bonder MJ, Stark AK, et al. Multitissue DNA methylation age predictor in mouse. Genome Biol. 18, 68.

83. Hogervorst E, Yaffe K, Richards M, Huppert FA. Hormone replacement therapy to maintain cognitive function in women with dementia. Cochrane Database Syst. Rev. 2009; 1: CD003799.

84. Gleason CE, Dowling NM, Wharton W, et al. Effects of hormone therapy on cognition and mood in recently postmenopausal women: findings from the randomized, controlled KEEPS-Cognitive and Affective study. PLoS Med. 2015; 12: e1001833.

85. Schneider L. Alzheimer’s disease and other dementias: update on research. Lancet Neurol. 2017; 16: 4–5.

86. Whitmer RA, Quesenberry CP, Zhou J, Yaffe K. Timing of hormone therapy and dementia: the critical window theory revisited. Ann. Neurol. 2011; 69: 163–9.

87. Horvath S. DNA methylation age of human tissues and cell types. Genome Biol. 2013; 14: R115.

88. Berson A, Nativio R, Berger SL, Bonini NM. Epigenetic Regulation in Neurodegenerative Diseases. Trends Neurosci. 2018; 41: 587–598.

89. Ianov L, Kumar A, Foster TC. Epigenetic regulation of estrogen receptor α contributes to age-related differences in transcription across the hippocampal regions CA1 and CANeurobiol. Aging, 2017; 49: 79–85.

90. Farrell LL, Schoenebeck JJ, Wiener P, Clements DN, Summers KM. The challenges of pedigree dog health: approaches to combating inherited disease. Canine Genet. Epidemiol. 2015; 2: 3.

91. Sun T, Warrington NM, Rubin JB. Why does Jack, and not Jill, break his crown? Sex disparity in brain tumors. Biol. Sex. Differ. 2012; 3: 3.

92. Wen PY, Kesari S. Malignant gliomas in adults. N. Engl. J. Med. 2008; 359: 492–507.

93. Schoemaker MJ, Swerdlow AJ, Higgins CD, Wright AF, Jacobs PA. UK Clinical Cytogenetics Group, Cancer incidence in women with Turner syndrome in Great Britain: a national cohort study. Lancet Oncol. 2008; 9: 239–46.

94. Dumanski JP, Rasi C, Lönn M. Mutagenesis. Smoking is associated with mosaic loss of chromosome Y. Science, 2015; 347: 81–83.

95. Dumanski JP, Lambert JC, Rasi C, et al. Mosaic Loss of Chromosome Y in Blood Is Associated with Alzheimer Disease. Am. J. Hum. Genet. 2016; 98: 1208–1219.

96. Forsberg LA. Loss of chromosome Y (LOY) in blood cells is associated with increased risk for disease and mortality in aging men. Human Genet. 2017; 136: 657–663.

97. Sun T, Warrington NM, Luo J, et al. Sexually dimorphic RB inactivation underlies mesenchymal glioblastoma prevalence in males. J. Clin. Invest. 2014; 124: 4123–33.

98. Harcourt-Brown T, Campbell J, Warren-Smith C, Jeffery ND, Granger NP. Prevalence of Chiari-like malformations in clinically unaffected dogs. J. Vet. Intern. Med. 2015; 29: 231–237.

99. Driver CJ, Volk HA, Rusbridge C, Van Ham LM. An update on the pathogenesis of syringomyelia secondary to Chiari-like malformations in dogs. Vet. J. 2013; 198: 551–559.

100. MacKillop E. Magnetic resonance imaging of intracranial malformations in dogs and cats. Vet. Radiol. Ultrasound 2011; 52: S42–S51.

101. Laubner S, Ondreka N, Failing K, Kramer M, Schmidt MJ. Magnetic resonance imaging signs of high intraventricular pressure--comparison of findings in dogs with clinically relevant internal hydrocephalus and asymptomatic dogs with ventriculomegaly. BMC Vet. Res. 2015; 11: 181.

102. Ryan CT, Glass EN, Seiler G, Zwingenberger AL, Mai W. Magnetic resonance imaging findings associated with lateral cerebral ventriculomegaly in English bulldogs. Vet. Radiol. Ultrasound 2014; 55: 292–299.

103. Cerda-Gonzalez S, Olby NJ, Griffith EH. Dorsal compressive atlantoaxial bands and the craniocervical junction syndrome: association with clinical signs and syringomyelia in mature Cavalier King Charles Spaniels. J. Vet. Intern. Med. 2015; 29: 887–892.

104. Marino DJ, Loughin CA, Dewey CW, et al. Morphometric features of the craniocervical junction region in dogs with suspected Chiari-like malformation determined by combined use of magnetic resonance imaging and computed tomography. Am. J. Vet. Res. 2012; 73: 105–111.

105. Rzechorzek NM, Liuti T, Stalin C, Marioni-Henry K. Restored vision in a young dog following corticosteroid treatment of presumptive hypophysitis. BMC Vet. Res. 2017; 13: 63.

